# Similarities and differences between IL11 and IL11RA1 knockout mice for lung fibro-inflammation, fertility and craniosynostosis

**DOI:** 10.1101/2020.12.10.420695

**Authors:** Benjamin Ng, Anissa A. Widjaja, Sivakumar Viswanathan, Jinrui Dong, Sonia P. Chothani, Stella Lim, Shamini G. Shekeran, Jessie Tan, Sebastian Schafer, Stuart A. Cook

**Affiliations:** Cardiovascular and Metabolic Disorders Program, Duke-National University of Singapore Medical School, Singapore; National Heart Research Institute Singapore, National Heart Centre Singapore, Singapore; MRC-London Institute of Medical Sciences, Hammersmith Hospital Campus, London, UK; National Heart and Lung Institute, Imperial College, London, UK

## Abstract

Genetic loss of function (LOF) in *IL11RA* infers IL11 signaling as important for fertility, fibrosis, inflammation and craniosynostosis. The impact of genetic LOF in *IL11* has not been characterized. We generated IL11-knockout *(Il11^-/-^)* mice, which are born in normal Mendelian ratios, have normal hematological profiles and are protected from bleomycin-induced lung fibro-inflammation. Noticeably, baseline IL6 levels in the lungs of *Il11^-/-^* mice are lower than those of wild-type mice and are not induced by bleomycin damage, placing IL11 upstream of IL6. Lung fibroblasts from *Il11^-/-^* mice are resistant to pro-fibrotic stimulation and show evidence of reduced autocrine IL11 activity. *Il11^-/-^* female mice are infertile. Unlike *Il11ra1^-/-^* mice, *Il11^-/-^* mice do not have a craniosynostosis-like phenotype and exhibit mildly reduced body weights. These data highlight similarities and differences between LOF in *IL11* or *IL11RA* while establishing further the role of IL11 signaling in fibrosis and stromal inflammation.

## Introduction

Interleukin 11 (IL11) was originally described as a factor important for hematopoiesis, notably platelet production, but more recently found to drive fibro-inflammatory disorders^1^. IL11 is a member of the IL6 family of cytokines, which share the gp130 coreceptor, but while IL6 has been studied in very great detail with an armamentarium of genetic tools to dissect its function, IL11 remains poorly characterised^1^.

It is apparent from the published literature that the majority of our understanding of the biology associated with loss-of-function (LOF) in IL11 signaling is inferred from studies of *IL11RA* mutant humans or mice^1^. The field of lL11 biology has lacked a mouse genetic model specific for IL11 LOF, which represents a gap in our understanding. This is important as, in the case of the family member IL6, there are both similarities and differences between effects of LOF in the *IL6* cytokine as compared to LOF in its cognate receptor alpha chain *(IL6RA*)^2,3^. As such, it is possible that the phenotype of *IL11RA* LOF may not map precisely to IL11 function. Furthermore, studies of IL11RA1 LOF have been conducted in a single mouse strain and there are additional genes in the targeted locus, which is a potential shortcoming.

Based on the genetic studies of IL11RA mutants, IL11 signaling is thought important for a number of phenotypes. *Il11ra1*-deleted female mice are infertile^4^ and mutation in *Il11ra1* in the mouse is associated with incompletely penetrant snout displacement and tooth abnormalities, a craniosynostosis-like syndrome^5^. Several human studies have identified individuals with *IL11RA* mutations who have features of craniosynostosis, joint laxity, scoliosis and delayed tooth eruption^5–7^. Unlike mice, female humans with mutations in *IL11RA* appear fertile^8^. In keeping with this, at the level of the general population there is no negative selection against predicted loss-of-function mutations in *IL11RA*, suggesting such mutations are not detrimental for replicative capacity^9^.

Here, we report the generation of mice with germline deletion of *Il11* that we characterise at baseline and in the context of pro-fibrotic stimulation *in vitro* and *in vivo.* We report similarities and differences between the phenotypes of *Il11^-/-^* and *Il11ra1^-/-^* mice.

## Results

### Generation and gross anatomical characterization of *Il11*-knockout mice

Three separate transcripts of mouse *Il11* have been annotated (ENSMUSG00000004371) and using Crispr/Cas9, we deleted Exon 2 to 4 of the longest transcript (ENSMUST00000094892.11: Il11-201). This deletion causes a reading frame shift after the first two amino acids of IL11 resulting in a mutant 62 amino acids peptide that does not align to any known peptide sequences resulting in the inactivation of all known transcripts (**Fig. 1A**). *Il11*-knockout mice (*Il11^-/-^*) were generated on a C57BL/6J background and genotypes were determined by sequencing and PCR (**Fig. 1B**). To address whether the mutant alleles resulted in the loss of *Il11* RNA expression, we isolated total RNA from whole lung tissue from *Il11^-/-^* mice and did not observe any detectable expression of *Il11* RNA by RT-qPCR in *Il11*^-/-^ mutants (**Fig. 1C**). We observed a slight (5-7% lower) but statistically significant reduction in body weights of male and female *Il11^-/-^* mice (10-12 weeks old) as compared to age and gender matched wild-type controls (**Fig. 1D**), which has not been reported in *Il11ra1^-/-^* mice of a similar age.

**Figure 1.**
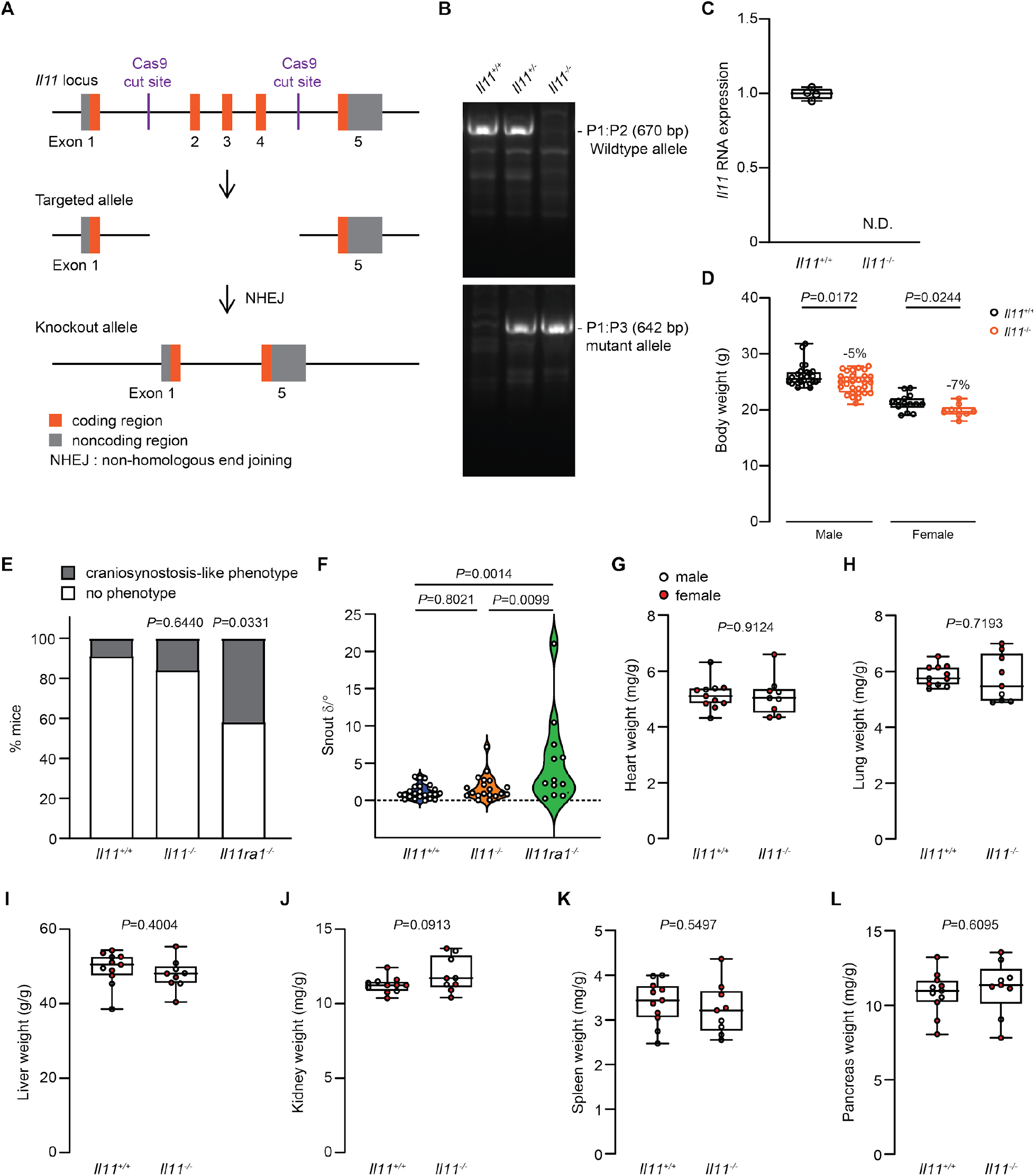
Generation and anatomical characterization of *Il11*-knockout mice. (**A**) Schematic design of Crispr/Cas9 mediated deletion of exons 2 to 4 of the mouse *Il11* locus. (**B**) Representative genotyping of *Il11*-deficient or wild-type mice, showing a wild-type band (670 bp) and mutant band (642 bp). (**C**) RT-qPCR of *Il11* expression in whole lung tissue from wild-type and *Il11^-/-^* mice (*n* =4). N.D., not detected. (**D**) Body weight of 10-12 week old wild-type (male *n*=26; female *n*=14) and *Il11^-/-^* mice (male *n*=29; female *n=9).* (**E**) Proportion of mice with craniosynostosis-like phenotype and (**F**) the degree of deviation from linear snout growth (δ/°) was determined in *Il11^-/-^* mice as compared to wild-type and *Il11ra1^-/-^* mice *(Il11^+/+^ n=23, Il11^-/-^ n*=19, *Il11ra1^-/-^ n*=12). (**G**) Heart weight-, (**H**) lung weight-, (**I**) liver weight-, (**J**) kidney weight-, (**K**) spleen weight- and (**L**) pancreas weight-to-bodyweight indices of male and female *Il11^-/-^* and wild-type mice (10-14 weeks of age). Data shown in C, D, G-L as: centre line, median value; box edges, 25th and 75th percentiles; whiskers, minimum and maximum values; E are shown as stacked bar graphs; F shown as violin plot. *P* values were determined by Fisher’s exact test in panel E, ANOVA (Tukey’s test) in F and Student’s *t*-test in D, G-L.

It is reported that approximately 40-50% of *Il11ra1^-/-^* mice display twisted snouts, a craniosynostosis-like phenotype^10^. We assessed the skulls of adult *Il11^-/-^* (*n*=19) and wild-type (*n*=23) mice (>12 weeks of age) and *Il11ra1^-/-^* mice (*n*=12), all on the same C57BL/6J background. We observed macroscopic snout deformities in 42% of *Il11ra1^-/-^* mice (5 out of 12 mice), similar to the published incidence^10^. Unlike *Il11ra1^-/-^* mice, we did not observe significant differences in the proportions of *Il11^-/-^* mice with snout deformities as compared to wild-type controls (*P*=0.64) (**Fig. 1E**). Further analysis revealed that there was no significant difference in the degree of sideward deviation of snout growth in *Il11^-/-^* mice as compared to wild-type mice, unlike *Il11ra1^-/-^* mice that were deviated (**Fig. 1F**). In gross anatomy studies, the indexed organ-to-body weight ratios of the heart, lung, liver, kidney, spleen and pancreas were comparable in *Il11^-/-^* and wild-type mice (**Fig. 1G-L**).

### Female *Il11*-knockout mice are infertile

Deletion of *Il11ra1* in mice leads to female infertility due to defective embryo implantation related to abnormal placental decidualization, whereas *Il11ra1^-/-^* males are fertile^4,11,12^. Intercrosses of heterozygotes *(Il11^+/-^)* gave rise to viable and apparently normal homozygous mutant (*Il11^-/-^*) mice in the expected Mendelian ratios (**Fig. 2A**). We determine whether maternal *Il11* expression is required for fertility by mating homozygous (*Il11^-/-^*) female mice with male mice of variable *IL11* genotype (*Il11^+/+^, Il11^+/-^* or *Il11^-/-^)* and found that female mice deficient for *Il11* never had a detectable pregnancy nor gave birth to offspring (**Fig. 2B**), which mirrors the infertility phenotype of homozygous *Il11ra1^-/-^* female mice ^4^ Crossing homozygous (*Il11^-/-^* male mice with either wild-type *(Il11^+/+^)* or heterozygous *(Il11^+/-^)* female mice resulted in viable offspring of expected Mendelian ratios. However, litter sizes derived from *Il11^-/-^* male mice were significantly smaller as compared to intercrosses of heterozygotes (**Fig. 2C**). Hence, similar to germline loss of *Il11ra1*, the loss of *Il11* expression results in female infertility and appears to affect male fertility, directly or indirectly, in mice.

**Figure 2.**
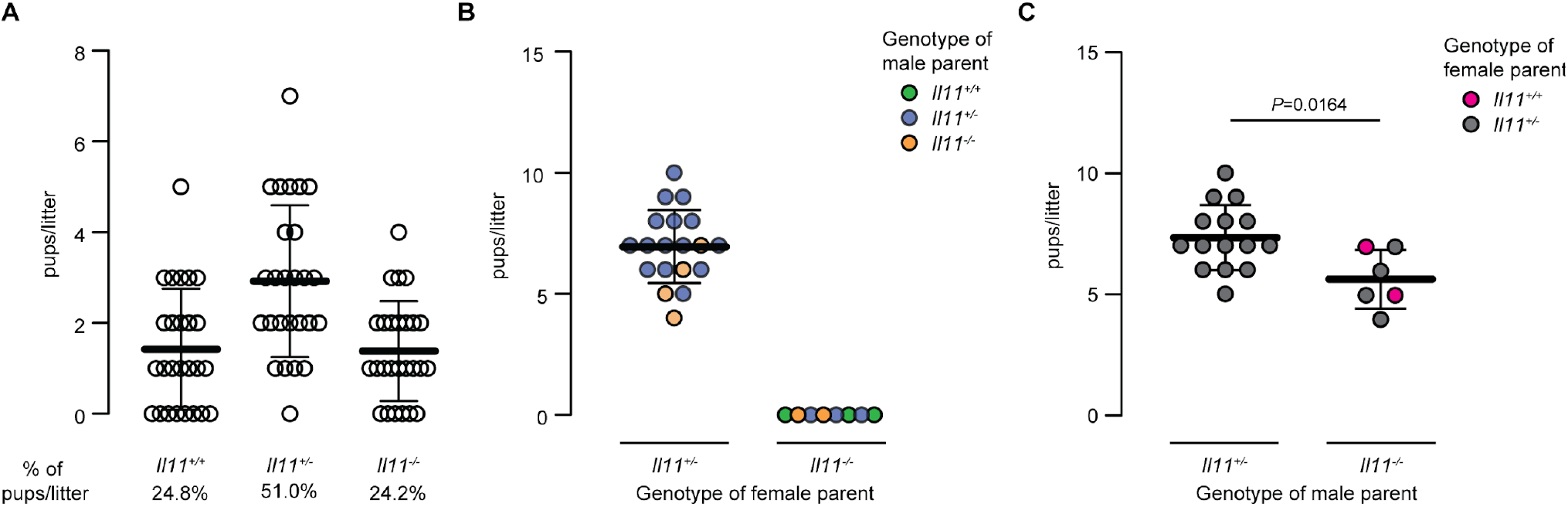
*Il11*-knockout female mice are infertile. (**A**) Genotype distribution of pups per litter in heterozygous breeding *(Il11^+/-^* X *Il11^+/-^; n*=26 litters). (**B**) Litter size based on parents’ genotype. (**C**) Litter size based on the breeding of heterozygous *Il11^+/-^* or homozygous *Il11^-/-^* male mice with wild-type or heterozygous *Il11^+/-^* female mice. Data shown as mean ± SD. *P* value was determined by Student’s *t* test.

### Blood hematology and chemistry profiles are normal in *Il11*-knockout mice

We evaluated the hematological profile of adult *Il11^-/-^* mice (10-14 weeks of age) and observed that null mice had normal peripheral red and white blood cell counts as well as normal platelet counts and volumes as compared to wild-type mice (**Table 1**). Likewise, we profiled serum chemistry and observed normal levels of serum markers of liver function (albumin, alanine aminotransferase, total bilirubin), kidney function (blood urea nitrogen, sodium and potassium) and bone turnover (alkaline phosphatase, calcium and phosphate) in *Il11^-/-^* mice (**Table 1**). These data indicate that *Il11^-/-^* mice have normal blood hematological and chemistry profiles under normal physiological conditions.

**Table. 1.**
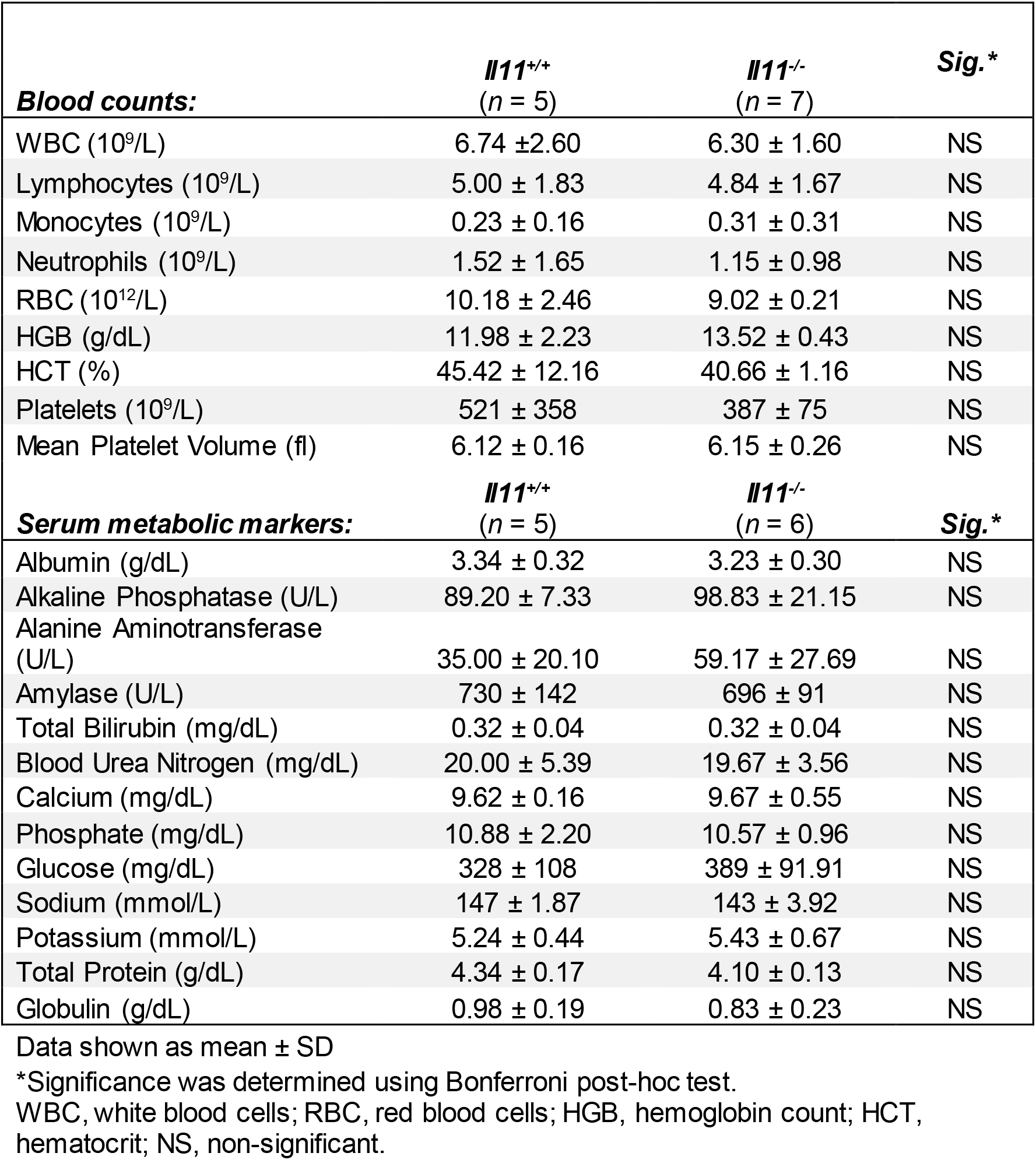
Hematology and serum metabolic profiles of *Il11^-/-^* mice.

| **Blood counts:** | ***Il11<sup>+/+</sup>***<br>**(n = 5)** | ***Il11<sup>-/-</sup>***<br>**(n = 7)** | **Sig.\*** |
| --- | --- | --- | --- |
| WBC (10<sup>9</sup>/L) | 6.74 ± 2.60 | 6.30 ± 1.60 | NS |
| Lymphocytes (10<sup>9</sup>/L) | 5.00 ± 1.83 | 4.84 ± 1.67 | NS |
| Monocytes (10<sup>9</sup>/L) | 0.23 ± 0.16 | 0.31 ± 0.31 | NS |
| Neutrophils (10<sup>9</sup>/L) | 1.52 ± 1.65 | 1.15 ± 0.98 | NS |
| RBC (10<sup>12</sup>/L) | 10.18 ± 2.46 | 9.02 ± 0.21 | NS |
| HGB (g/dL) | 11.98 ± 2.23 | 13.52 ± 0.43 | NS |
| HCT (%) | 45.42 ± 12.16 | 40.66 ± 1.16 | NS |
| Platelets (10<sup>9</sup>/L) | 521 ± 358 | 387 ± 75 | NS |
| Mean Platelet Volume (fl) | 6.12 ± 0.16 | 6.15 ± 0.26 | NS |
| **Serum metabolic markers:** | ***Il11<sup>+/+</sup>***<br>**(n = 5)** | ***Il11<sup>-/-</sup>***<br>**(n = 6)** | **Sig.\*** |
| Albumin (g/dL) | 3.34 ± 0.32 | 3.23 ± 0.30 | NS |
| Alkaline Phosphatase (U/L) | 89.20 ± 7.33 | 98.83 ± 21.15 | NS |
| Alanine Aminotransferase (U/L) | 35.00 ± 20.10 | 59.17 ± 27.69 | NS |
| Amylase (U/L) | 730 ± 142 | 696 ± 91 | NS |
| Total Bilirubin (mg/dL) | 0.32 ± 0.04 | 0.32 ± 0.04 | NS |
| Blood Urea Nitrogen (mg/dL) | 20.00 ± 5.39 | 19.67 ± 3.56 | NS |
| Calcium (mg/dL) | 9.62 ± 0.16 | 9.67 ± 0.55 | NS |
| Phosphate (mg/dL) | 10.88 ± 2.20 | 10.57 ± 0.96 | NS |
| Glucose (mg/dL) | 328 ± 108 | 389 ± 91.91 | NS |
| Sodium (mmol/L) | 147 ± 1.87 | 143 ± 3.92 | NS |
| Potassium (mmol/L) | 5.24 ± 0.44 | 5.43 ± 0.67 | NS |
| Total Protein (g/dL) | 4.34 ± 0.17 | 4.10 ± 0.13 | NS |
| Globulin (g/dL) | 0.98 ± 0.19 | 0.83 ± 0.23 | NS |

Data shown as mean ± SD
\*Significance was determined using Bonferroni post-hoc test.
WBC, white blood cells; RBC, red blood cells; HGB, hemoglobin count; HCT, hematocrit; NS, non-significant.

### IL11 is required for myofibroblast differentiation

We previously found that TGFβ1-induced myofibroblast transdifferentiation was impaired in *Il11ra1^-/-^* lung fibroblasts^13^. To examine whether the loss of the endogenous IL11 autocrine feed-forward loop similarly perturbed fibroblast activation, we stimulated lung fibroblasts from *Il11^-/-^* mice with recombinant mouse TGFβ1 or IL11 (5 ng/ml; 24 hours) and monitored fibroblast activation using automated high-throughput immunofluorescence imaging and Sirius red-based quantification of secreted collagen. In keeping with the data from *Il11ra1*-deleted fibroblasts, the differentiation of *Il11^-/-^* fibroblasts into ACTA2^+ve^ and COL1A1 expressing myofibroblasts following TGFβ1 stimulation was significantly diminished (**Fig. 3A-B**). Cell proliferation (as determined by EdU^+ve^ staining) and secreted collagen levels into the culture supernatant were also significantly reduced in *Il11^-/-^* fibroblasts following TGFβ1 stimulation (**Fig. 3C-D**).

**Figure 3.**
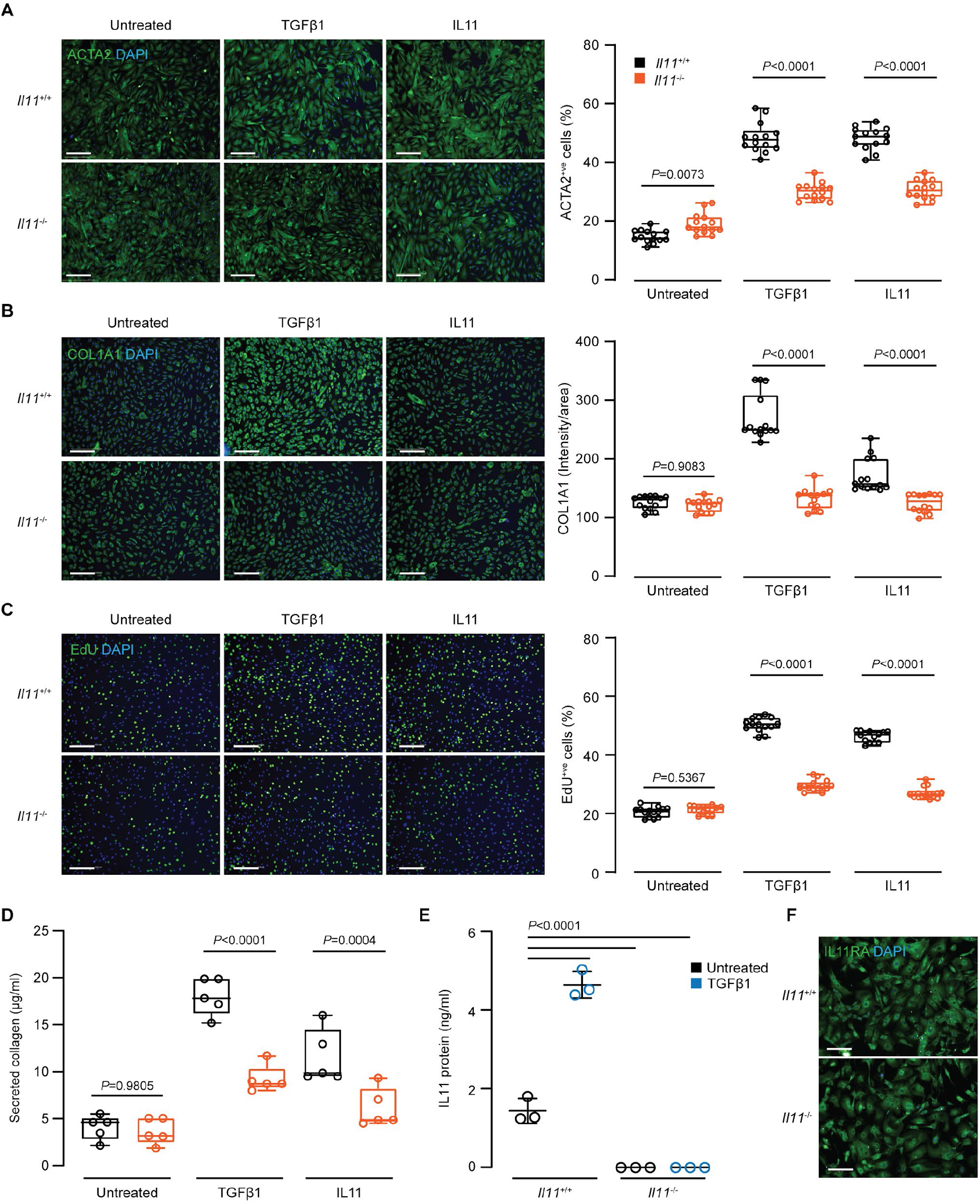
Reduced activation of primary lung fibroblasts from *Il11*-knockout mice. Automated immunofluorescence quantification of (**A**) ACTA2^+ve^ cells, (**B**) COL1A1 expression (intensity/area) and (**C**) EdU^+ve^ cells in TGFβ1- or IL11-treated lung fibroblasts from *Il11^-/-^* or wild-type mice (5 ng/ml, 24h). One representative dataset from two independent biological experiments is shown (14 measurements per condition per experiment). (**D**) Secreted collagen concentrations in supernatant of cells treated as described in A-B were quantified (*n*=5). (**E**) Secreted IL11 in culture supernatant from TGFβ1-treated lung fibroblasts from *Il11^-/-^* or wild-type mice (5 ng/ml, 24h; *n*=3). (**F**) Immunofluorescence images of IL11RA expression in lung fibroblasts from *Il11^-/-^* or wild-type mice. Scale bars: 200 μm. Data in A-D are shown as: centre line, median value; box edges, 25th and 75th percentiles; whiskers, minimum and maximum values; and shown as mean ± SD in E. *P* values in A-D were determined by ANOVA (Sidak’s test) and by ANOVA (Tukey’s test) in E.

We next addressed whether disruption of the IL11 locus prevented IL11 protein expression by performing ELISA on culture supernatants and found that IL11 protein was not expressed by *Il11^-/-^* lung fibroblasts at baseline or after TGFβ1 stimulation (**Fig. 3E**). Interestingly, recombinant mouse IL11 (5ng/ml) did not fully restore pro-fibrotic phenotypes in *Il11^-/-^* fibroblasts (**Fig. 3A-D**). We assessed the expression of IL11RA in *Il11^-/-^* lung fibroblasts by immunofluorescence staining and detected comparable levels of IL11RA expression between *Il11^-/-^* and wild-type cells (**Fig. 3F**). This suggests that the autocrine loop of IL11 in the local environs of the cell is of greater importance for pro-fibrotic activity than exogenous IL11.

### Bleomycin-induced pulmonary fibrosis is attenuated in *Il11*-knockout mice

We recently showed that IL11 expression is elevated in the lung after bleomycin (BLM)- induced injury and that BLM-induced lung fibrosis is attenuated in *Il11ra1^-/-^* mice^14^. To determine if the genetic deletion of *Il11* provided similar protection, we challenged *Il11^-/-^* mice with a single dose of BLM oropharyngeally, and assessed the lungs 14 days thereafter (**Fig. 4A**). By gross morphology analysis, we observed reduced macroscopic lung damage in *Il11^-/-^* mice than that seen in wild-type mice (**Fig. 4B**). Consistent with this, blinded histological analysis of Masson’s trichrome stained lung sections showed that *Il11^-/-^* mice had reduced parenchymal disruption and fibrosis (**Fig. 4C-D**). These changes were associated with significantly lower total lung hydroxyproline (collagen) content in *Il11^-/-^* mice (**Fig 4E**).

**Figure 4.**
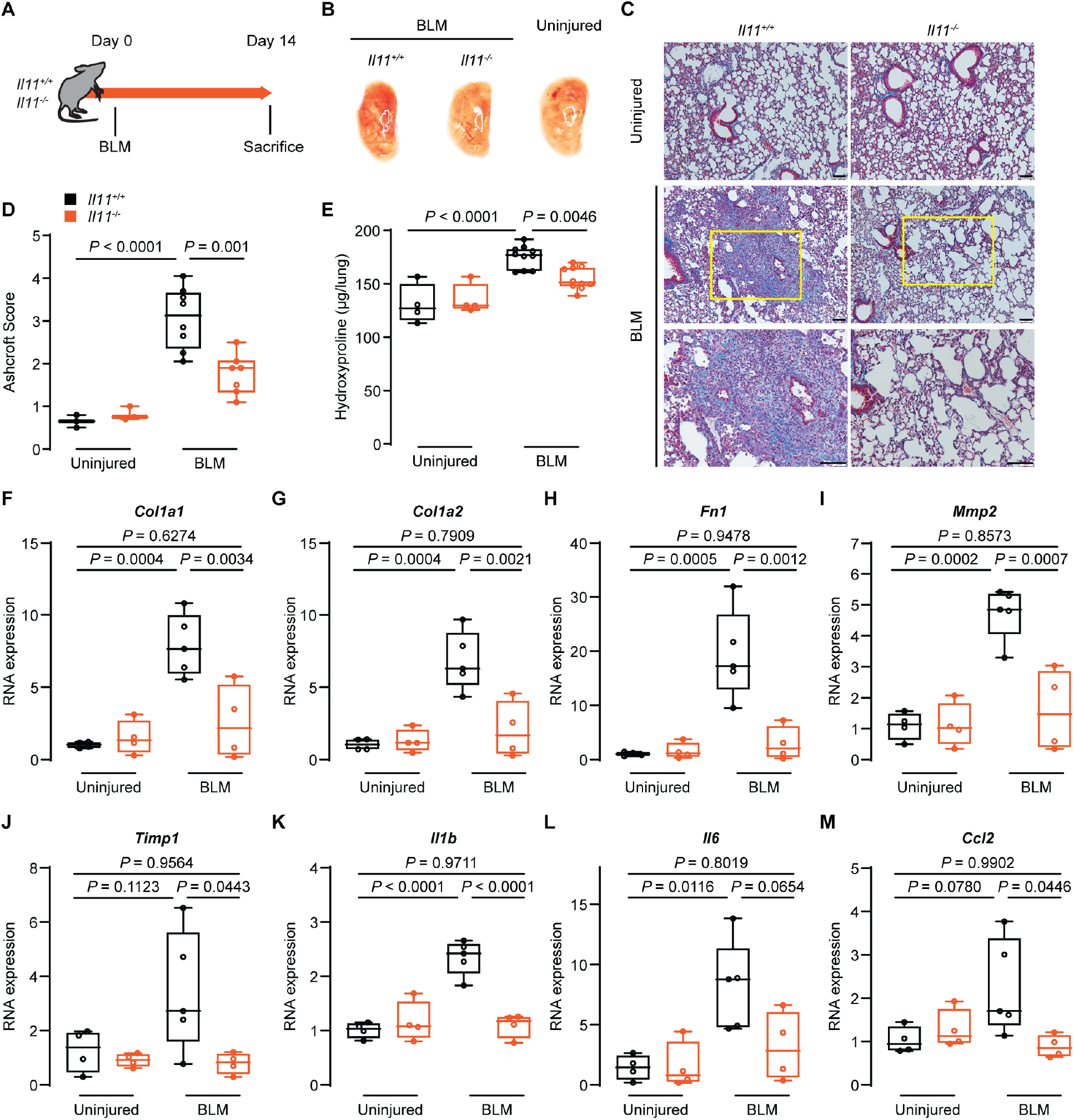
Bleomycin-induced pulmonary fibrosis is attenuated in *Il11*-knockout mice. (**A**) Schematic showing the induction of lung fibrosis in *Il11^-/-^* and wild-type mice. A single dose of bleomycin (BLM) was administered oropharyngeally and the mice were sacrificed 14 days post-BLM. (**B**) Representative gross lung anatomy of *Il11^-/-^* and wild-type mice 14 days post-BLM. (**C**) Masson’s trichrome staining of lung sections, (**D**) histology assessment of fibrosis and (**E**) total lung hydroxy proline content of *Il11^-/-^* and wildtype mice 14 days post-BLM. Scale bars: 100 μm. Relative RNA expression of (**F**) *Col1a1*, (**G**) *Col1a2*, (**H**) *Fn1*, (**I**) *Mmp2*, (**J**) *Timp1*, (**K**) *Il1b*, (**L**) *Il6* and (**M**) *Ccl2* in lung lysates from *Il11^-/-^* and wild-type mice 14 days post-BLM (*n*=4). Data shown as: centre line, median value; box edges, 25th and 75th percentiles; whiskers, minimum and maximum values. *P* values were determined by ANOVA (Tukey’s test).

Evaluation of fibrotic gene expression in lung lysates showed reduced RNA levels of extracellular matrix and protease genes such as *Col1a1, Col1a2, Fn1, Mmp2* and *Timp1* in BLM-challenged *Il11^-/-^* mice as compared to wild-type mice (**Fig. 4F-J**). Reduced expression of several inflammatory response genes (such as *Il1b, Il6* and *Ccl2)* were also seen in the lungs of *Il11^-/-^* mice following BLM injury (**Fig. 4K-M**).

Western blot analysis showed that IL11 protein expression was strongly upregulated in the lungs of BLM-injured wild-type mice and was not expressed at all in the lungs of *Il11^-/-^* mice at all, as expected (**Fig. 5A**). Furthermore, in BLM-treated *Il11^-/-^* mice, lung protection was accompanied by reduced pulmonary fibronectin and IL6 protein expression (**Fig. 5A**) and reduced ERK activation (**Fig. 5B**). Notably, lung IL6 levels in *Il11^-/-^* mice were lower than wild type control levels in the absence of lung injury. These data show that *Il11^-/-^* mice are protected from BLM-induced lung fibrosis and inflammation, similar to *Il11ra1^-/-^* mice, while confirming the importance of IL11- stimulated ERK activation for these phenotypes^14,15^.

**Figure 5.**
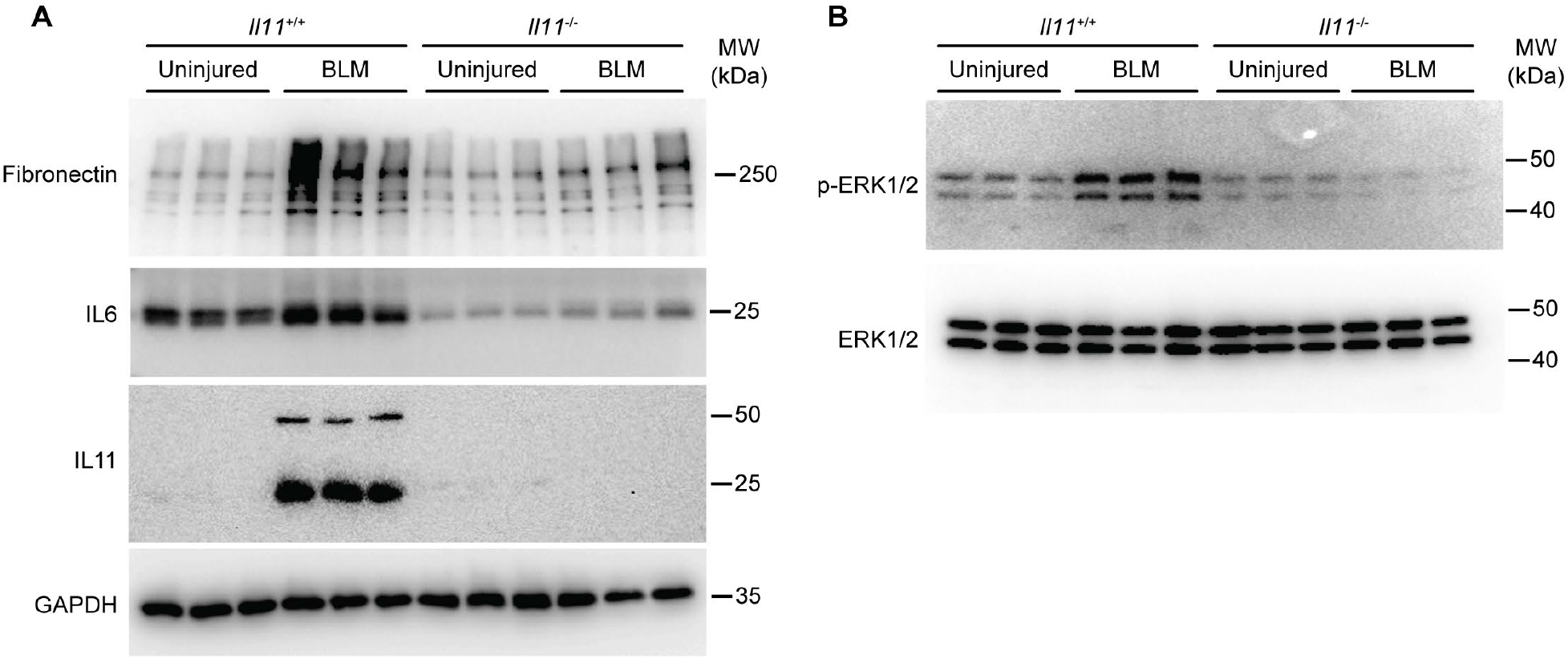
Bleomycin-induced pulmonary fibronectin, IL6 and IL11 expression and ERK activation in *Il11*-knockout and wild type mice. Western blots of (**A**) fibronectin, IL6 and IL11 and (**B**) phosphorylated and total ERK1/2 in lung homogenates of *Il11^-/-^* and wild-type mice 14 days post-BLM (*n*=3 for each genotype/condition). Uncropped blot images are shown in **Supplementary Figure 1**.

## Discussion

Here we provide a phenotypic description of mice with germline deletion of *Il11* and explore how this relates to the phenotypes of *Il11ra1-null* mice. We show that IL11 signaling in *Il11^-/-^* mice is important for fibrotic phenotypes in fibroblasts *in vitro* and lung fibrosis *in vivo.* These data, taken together with recent studies of *Il11ra1^-/-^* mice and the use of anti-IL11 or anti-IL11RA antibodies in mouse models of fibrosis^13,14,16^, firmly establish IL11 signaling as of central importance for fibrosis.

*Il11ra1^-/-^* mice have normal hematopoiesis at baseline and after bone marrow or hemolytic stress^11^ and long term use of neutralizing anti-IL11 or anti-IL11RA antibodies have no effect on blood counts in mice^16^. In agreement with mouse data, there is no description of hematological abnormalities in patients with *Il11ra1* mutations^17^. While IL11 is still considered by some to have a role in hematopoiesis, the data shown here in *Il11^-/-^* mice provide a further line of evidence that IL11 is unrelated to this biology, although we did not study bone marrow stress.

It was surprising to observe differences in skull morphology between *Il11^-/-^* and *Il11ra1^-/-^* genotypes. These differences are unlikely to be accounted for by genetic factors as both strains are maintained on C57Bl6/J backgrounds. The fact that *Il11^-/-^* mice do not have snout deformities implies that this is unrelated to the loss of IL11 signaling *perse.* However, a biallelic non-synonymous variant in *IL6ST*, which results in a selective loss of IL11-signaling, conferred a craniosynostosis phenotype, which replicated in a mouse model^18^. Intriguingly, while LOF mutations in *IL11* are common in the general population they have not been associated with craniosynostosis despite large scale sequencing projects^7^ whereas *IL11RA* mutations are widely reported ^5,6^. These data suggest the effect of IL11 signaling on craniosynostosis phenotypes is nuanced and/or differs depending on whether *IL11RA* or *IL11* is disrupted.

There are parallels between the variant phenotypes between *Il11ra1*- and *Il11*- deficient mice and those seen for *Il6-* and *Il6ra*-deficient mice. For instance, physiologically important immune phenotypes associated with loss of canonical IL6-mediated JAK/STAT signaling are seen with both *Il6*- or *Il6ra* genotypes. However, dissimilar ERK-mediated wound healing phenotypes are only seen in *Il6ra* mutants, which are dominant over *Il6* LOF effects (i.e. IL6-independent)^3^. Furthermore, *Il6* or *Il6ra* mutant mice also have different responses to experimental models of colitis^2^.

A possible explanation for the difference between alpha chain and ligand mutants could relate to reduced/no interaction of mutant alpha chains with the gp130. Thus, LOF in one alpha chain (e.g. IL6R) might reduce its competition for the shared gp130 receptor and potentiate the activity of another gp130-binding alpha chain (e.g. IL11RA). This premise could account for the increased ERK signaling seen in IL6RA mutant mice, perhaps reflecting increased IL11-driven ERK signaling^3^. The idea of diminished interaction of mutant alpha chains with gp130 is suggested further by human genetics: autosomal recessive mutations in gp130 are associated with craniosynostosis, whereas autosomal dominant variation is not^19,20^.

There is variation in reported fertility phenotypes associated with LOF mutations in the IL11 pathway in mice and humans (**Table 2**). Female mice lacking *Il11ra1* are infertile due to defective decidualization in response to embryo implantation ^4,12^. In contrast, women with homozygous *IL11RA* variants appear able to reproduce ^8^ which could be explained by species differences in decidualization^21^. In this study, we found that *Il11^-/-^* female mice are infertile, consistent with a recent report ^22^. Interestingly, we observed a reduction in litter sizes from *Il11^-/-^* male mice, suggesting additional male-related effects of IL11 on fertility.

**Table. 2.** Fertility reported with genetic loss-of-function in IL11 signaling in mice and humans.

| Gene | Mutation/variant | Gender effects | Reference |
| --- | --- | --- | --- |
| **Human** |  |  |  |
| *IL11RA* | Exon 4 donor splice site (c.479+6T>G) | Females: fertile | (6) |
| **Mouse** |  |  |  |
| *Il6st* | Gp130 p.R279Q (selective IL11 signaling deficiency) | Females: homozygous mutants have reduced litter size<br>Males: homozygous mutants have reduced litter size | (18) |
| *Il11ra1* | Deletion of Exons 8 to 13 | Females: homozygous mutants are infertile due to defective decidulization<br>Males: normal fertility | (4,11) |
| *Il11ra1* | Hypomorphic mutations | Females: fertility of homozygous mutants are severely impaired<br>Males: normal fertility | (12) |
| *Il11* | Crispr/Cas9 frameshift in Exon 3 | Females: homozygous mutants are infertile (data not shown)<br>Males: not reported | (22) |
| *Il11* | Crispr/Cas9 deletion of Exons 2 to 4. Frameshift from 2<sup>nd</sup> amino acid (aa), mutant 62 aa protein. | Females: homozygous mutants are infertile<br>Males: homozygous mutants have reduced litter size | Ng. et al. 2020 (This study) |

While IL11 is increasingly recognized as important for tissue fibrosis, more recent data has shown a role for IL11 signaling in inflammatory fibroblasts and stromal immunity in the lung and colon^23 15^ Fitting with this, we found that *Il11^-/-^* mice were protected from BLM-induced lung inflammation with lower *Il1b, Il6* and *ccl2* mRNA levels. More strikingly, at the protein level, IL6 was not only not upregulated in the injured lung of *Il11^-/-^* mice but IL6 levels were also lower in *Il11^-/-^* mouse lungs at baseline.

In conclusion, loss of IL11 signaling due to mutation in either *IL11* or *IL11RA* is protective against fibrosis that relates, in part, to reduced autocrine IL11 activity in myofibroblasts^13^. However, the craniosynostosis phenotypes seen in *IL11RA* deficient humans and mice are not apparent in *Il11^-/-^* mice, suggesting that this may not be directly due to defective IL11 signaling. While IL6 is one of the most studied genes in the literature, increasing evidence supports a role for IL11 upstream of IL6 that has potential large implications. To facilitate further analysis of IL11 LOF the *Il11^-/-^* mice described in this manuscript have been made available to the scientific community at the Jackson Laboratories repository.

## Materials and Methods

### Animal studies

All experimental procedures were approved and conducted in accordance with guidelines set by the Institutional Animal Care and Use Committee at SingHealth (Singapore) and the SingHealth Institutional Biosafety Committee and with the recommendations in the *Guidelines on the Care and Use of Animals for Scientific Purposes of the National Advisory Committee for Laboratory Animal Research* (NACLAR). Animals were maintained in a specific pathogen-free environment and given ad libitum access to food and water.

#### Generation of Il11-knockout mice

Crispr/Cas9 technique was used to knock out the IL11 gene (ENSMUST00000094892.11: Il11-201 transcript). Specific single guide RNA (sgRNA) sequences with recognition sites on introns 1 and 4 along with Cas9 were microinjected into fertilized C57BL/6J zygotes, and subsequently transferred into pseudopregnant mice (Shanghai Model Organisms, Centre, Inc). It is predicted that this deletion would cause a shift in the reading frame from the splicing of coding sequences present within exons 1 and 5, resulting in the generation of a mutant peptide (62 amino acids in length) that does not align to that of known proteins. This effectively results in the inactivation of the gene. Successfully generated F0 mice were identified by PCR and sequencing and further backcrossed to wild type C57BL/6J mice and maintained on this background. Wild-type allele was identified by a 670 bp PCR product using genotyping primer sequences as follows: P1: 5′-CGGGGGCGGACGGGAGACG-3′ and P2: 5′-CCAGGAGGGATCGGGTTAGGAGAA-3′. Whereas, mutant (knockout) alleles were identified by a second PCR reaction using primers P1 and P3: 5′- CAGCTAGGGACGACACTTGAGAT-3′.

#### Bleomycin model of lung injury

The bleomycin model of lung fibrosis was performed as previously described^14^. Briefly, female mice (8-10 weeks of age) were anesthetized by isoflurane inhalation and subsequently administered a single dose of bleomycin (Sigma-Aldrich) oropharyngeally at 0.5 mg/kg body weight in a volume of saline not exceeding 50 μl per mouse. Uninjured animals received equal volumes of saline as sham controls. Mice were sacrificed 14 days post-bleomycin administration and lungs were collected for downstream analysis.

### Analysis of craniosynostosis-like snout phenotype

*Il11ra1^-/-^* mice on a C57BL/6J background were originally described in ^11^ and obtained from The Jackson’s laboratory. For the analysis of skull phenotypes of *Ill1^-/-^, Il11ra1^-/-^* and wild-type mice, the heads of both male and female adult mice (>12 weeks of age) were dissected and cleaned by removing the soft tissue surrounding the skull. The bones were fixed in 10% neutral buffered formalin. Classification was performed by visual scoring of the presence (craniosynostosis-like phenotype) or absence (no phenotype) of twisted snouts by two independent investigators blinded to genotypes, and concordant scores were obtained between the investigators. Deviation from linear nasal bone growth was determined by assessing the angle between the tip of the snout and the sagittal suture at the base of the frontal bone by ImageJ software analysis (v1.8).

### Hematologic analysis

Blood was collected by cardiac puncture from anesthetized male and female adult mice (10-14 weeks of age). Differential red and white cell counts, hematocrit, hemoglobin and platelet counts were determined using the VetScan HM5 hematology analyzer (Abraxis Inc.). Blood chemistry profiles from male and female mice were determined using the VetScan VS2 system, partnered with the VetScan comprehensive diagnostic profile discs (Abraxis Inc.).

### Reagents

Recombinant proteins: Mouse TGFβ1 (7666-MB, R&D Systems), mouse IL11 (rmIL11, UniProtKB: P47873, GenScript). Antibodies: anti-smooth muscle actin (ab7817, abcam), anti-Collagen I (ab34710, abcam), anti-IL11RA (ab125015, abcam), goat anti-mouse Alexa Flour 488-conjugated secondary antibody (ab150113, abcam), goat anti-rabbit Alexa Flour 488-conjugated secondary antibody (ab150077, abcam), DAPI (D1306, Thermo Fisher Scientific). Primary antibodies for Western blots include: anti-Fibronectin antibody (ab2413, Abcam); anti-IL6 (12912, Cell Signaling); anti-p-ERK1/2 (4370, Cell Signaling), anti-ERK1/2s (4695, Cell Signaling) and anti-GAPDH (2118, Cell Signaling). Monoclonal anti-IL11 antibody (X203), generated in our previous study^14^, was used to detect IL11 protein expression in tissue lysates. Secondary antibodies for Western blots include: anti-rabbit HRP (7074, Cell Signaling) or antimouse HRP (7076, Cell Signaling).

### Primary mouse lung fibroblasts cultures

Primary mouse lung fibroblasts were isolated from 8-12 weeks old *Il11^+/+^* and wild-type mice as previously described^14^. Tissues were minced, digested for 30 minutes with mild agitation at 37°C in DMEM (11995-065, Gibco) containing 1% penicillin/streptomycin (P/S, 15140-122, Gibco) and 0.14 Wunsch U ml^-1^ Liberase (5401119001, Roche). Cells were subsequently cultured in complete DMEM supplemented with 10% fetal bovine serum (10500, Hyclone), 1% P/S, in a humidified atmosphere at 37 °C and 5% CO2. Fresh medium was renewed every 2-3 days. Fibroblasts were allowed to explant from the digested tissues and enriched via negative selection with magnetic beads against mouse CD45 (leukocytes), CD31 (endothelial) and CD326 (epithelial) using a QuadroMACS separator (Miltenyi Biotec) according to the manufacturer’s protocol. All experiments were carried out at low cell passage (<P3) and cells were cultured in serum-free media for 16 hours prior to stimulation.

### Operetta high-content imaging and analysis

Immunofluorescence imaging and quantification of fibroblast activation were performed on the Operetta High Content Imaging System (PerkinElmer) as previously described^13,14^. Briefly, lung fibroblasts were seeded in 96-well CellCarrier black plates (PerkinElmer) and following experimental conditions, the cells were fixed in 4% paraformaldehyde (Thermo Fisher Scientific) and permeabilized with 0.1% Triton X-100 in phosphate-buffered saline (PBS). EdU-Alexa Fluor 488 was incorporated using Click-iT EdU Labelling kit (C10350, Thermo Fisher Scientific) according to manufacturer’s protocol. The cells were then incubated with primary antibodies (anti-ACTA2 or anti-COL1A1) and visualized using anti-mouse or anti-rabbit Alexa Flour 488-conjugated secondary antibodies. Plates were scanned and images were collected with the Operetta high-content imaging system (PerkinElmer). Each treatment condition was run in duplicate wells, and 14 fixed fields were imaged and analysed per condition. The percentage of activated myofibroblasts (ACTA2^+ve^ cells) and proliferating cells (EdU^+ve^ cells) was quantified using the Harmony software version 3.5.2 (PerkinElmer). Quantification of COL1A1 immunostaining was performed using the Columbus software (version 2.7.2, PerkinElmer), and fluorescence intensity was normalized to cell area.

### Lung histology analysis

Freshly dissected lungs from *Il1^-/-^* and wild-type mice were fixed in 10% formalin overnight, dehydrated and embedded in paraffin and sectioned for Masson’s trichrome staining as described previously^14^. Histological analysis for fibrosis was performed blinded to genotype and treatment exposure according to Ashcroft scoring method^24^.

### Quantification of collagen content in culture supernatant and lung tissue

Detection of soluble collagen in the supernatant of lung fibroblasts cultures were performed as previously described^14^. Briefly, the cell culture supernatant was concentrated using a Polyethylene glycol concentrating solution (90626, Chondrex) and collagen content was quantified using a Sirius red collagen detection kit (9062, Chondrex), according to the manufacturer’s protocol. Total lung hydroxyproline content in the right lobes of mice were measured as previously described^14^, using the Quickzyme Total Collagen assay kit (Quickzyme Biosciences).

### ELISA

Detection of secreted IL11 into the supernatant of lung fibroblast cultures was performed using the mouse IL-11 DuoSet ELISA kit according to manufacturers’ instructions.

### RT-qPCR

Total RNA was extracted from snap-frozen mouse lung tissues using Trizol reagent (Invitrogen) followed by RNeasy column (Qiagen) purification and cDNA was prepared using an iScript cDNA synthesis kit (Biorad) following manufacturer’s instructions. Quantitative RT-PCR gene expression analysis was performed with QuantiFast SYBR Green PCR kit (Qiagen) using a StepOnePlus (Applied Biosystem). Relative expression data were normalized to *Gapdh* mRNA expression using the 2^-ΔΔCt^ method. Primers sequences used for are as follows: *Il11* 5′- AATTCCCAGCTGACGGAGATCACA-3′ and 5′-TCTACTCGAAGCCTTGTCAGCACA-3′; *Col1a1* 5′-GGGGCAAGACAGTCATCGAA-3′ and 5′-GTCCGAATTCCTGGTCTGGG-3′; *Col1a2* 5′-AGGATTGGTCAGAGCAGTGT-3′ and 5′-TCCACAACAGGTGTCAGGGT-3′; *Fn1* 5′-CACCCGTGAAGAATGAAGA-3′ and 5′-GGCAGGAGATTTGTTAGGA-3′; *Mmp2* 5′-ACAAGTGGTCCGCGTAAAGT-3′ and 5′-AAACAAGGCTTCATGGGGGC-3′; *Timp1* 5′-GGGCTAAATTCATGGGTTCC-3′ and 5′-CTGGGACTTGTGGGCATATC-3′; *Ccl2* 5′-GAAGGAATGGGTCCAGACAT-3′ and 5′-ACGGGTCAACTTCACATTCA-3′; *Il6* 5′-CTCTGGGAAATCGTGGAAAT-3′ and 5′-CCAGTTTGGTAGCATCCATC-3′; Il1b 5′-CACAGCAGCACATCAACAAG-3′ and 5′- GTGCTCATGTCCTCATCCTG-3′; *Gapdh* 5′- CTGGAAAGCTGTGGCGTGAT-3 and 5′- GACGGACACATTGGGGGTAG-3′.

### Western blot analysis

Total proteins were extracted from snap-frozen mouse lung tissues using RIPA lysis buffer (Thermo Fisher Scientific) and separated by SDS-PAGE, transferred to a PVDF membrane (Biorad), and incubated overnight with the appropriate primary antibodies. Blots were visualized using the ECL detection system (Pierce) with the appropriate secondary antibodies.

### Statistical analysis

All statistical analyses were performed using Graphpad Prism (version 8). Statistical analyses were performed using two-sided Student’s t-test, or ANOVA as indicated in the figure legends. For comparisons between multiple treatment groups, *P* values were corrected for multiple hypothesis testing using Sidak’s test, Tukey’s test or Bonferonni’s post-hoc test. *P* values <0.05 were considered statistically significant.

## Data availability

All data generated and analysed in the current study are presented in the manuscript or available from the corresponding author upon request.

## Acknowledgements

We would like to acknowledge N. Ko and B. L. George for their support. This research is supported by the National Medical Research Council (NMRC), Singapore STaR awards (NMRC/STaR/0029/2017), MOH-CIRG18nov-0002, Goh Foundation, Tanoto Foundation to S.A.C. A.A.W. is supported by NMRC/OFYIRG/0053/2017. S.S is supported by NMRC/OFYIRG18nov-0014.

## Author Contributions

B.N., A.A.W. and S.A.C. designed the study. B.N., A.A.W., S.V., J.D., S.L., S.G.S. and J.T. performed experiments. B.N., A.A.W., S.V. and S.P.C. analyzed the data. B.N., A.A.W., S.S. and S.A.C. prepared and wrote the manuscript with input from the other co-authors.

